# Metabolic enhancement of mammalian developmental pausing

**DOI:** 10.1101/2022.08.22.504730

**Authors:** Vera A. van der Weijden, Maximilian Stoetzel, Beatrix Fauler, Dhanur P. Iyer, Mohammed Shahraz, David Meierhofer, Steffen Rulands, Theodore Alexandrov, Thorsten Mielke, Aydan Bulut-Karslioglu

## Abstract

The quest to model and modulate embryonic development became a recent cornerstone of stem cell and developmental biology. Mammalian developmental timing is adjustable in vivo by preserving preimplantation embryos in a dormant state called diapause. Inhibition of the growth regulator mTOR (mTORi) pauses mouse development in vitro, yet constraints to pause duration are unrecognized. By comparing the response of embryonic and extraembryonic stem cells to mTORi-induced pausing, we identified lipid usage as a bottleneck to developmental pausing. Enhancing fatty acid oxidation (FAO) boosts embryo longevity, while blocking it reduces the pausing capacity. Genomic and metabolic analyses of single embryos point toward a deeper dormant state in FAO-enhanced pausing and reveal a link between lipid metabolism and embryo morphology. Our results lift a constraint on in vitro embryo survival and suggest that lipid metabolism may be a critical metabolic transition relevant for longevity and stem cell function across tissues.

**One-Sentence Summary:** Facilitating fatty acid oxidation by carnitine supplementation enhances mTOR inhibition-mediated developmental pausing.

## Introduction

Dormancy maintains cells in a reversible non-proliferative state and is a major component of stem cell function^1^. Although mainly explored in the context of adult stem cell activity, dormancy is also employed in early development as a means to preserve the non-implanted embryo under certain circumstances^2^. Perturbation of dormancy entry and exit leads to exhaustion of stem cell pools, compromised regeneration, and risks reproductive success in many mammals. Thus, it is of great importance to identify cellular mechanisms of establishment and maintenance of dormancy as well as exit from it.

The cellular niche, signaling, and metabolism are all involved in the active rewiring of cellular networks during the transition from proliferation to dormancy^3-9^. The mTOR pathway acts as a rheostat to adjust cellular growth and anabolism according to environmental and metabolic signals. Due to this prominent role in growth control, the mTOR pathway is also a central regulator of dormancy across tissues, and its hyperactivation tilts the dormancy-activation balance leading to stem cell exhaustion^10-12^. Previously, we showed that inhibition of mTOR (mTORi) induces a diapause-like dormant state in vitro in embryonic stem cells (ESCs) and mouse preimplantation embryos ^13^. Mouse blastocyst-stage embryos can be reliably maintained in mTORi-induced diapause for a few weeks, however significant embryo loss occurs over time^13^. Importantly, the inner cell mass (ICM) and trophectoderm (TE) show distinct responses upon induction of dormancy in vitro through mTORi or in vivo through hormonal diapause^13^. Namely, the ICM and TE show independent dynamics of global histone acetylation, genomic activity, and rate of proliferation in response to the same conditions inducing diapause^13-15^.

Although dormancy as an end-state has common characteristics like loss of proliferation and global reduction in metabolic activity, how different cells respond to dormancy cues and the specific steps taken on this route are unclear. This is largely due to inaccessibility and limited numbers of dormant cells in vivo, and their tendency to spontaneously activate upon isolation from their niche^16^. Here, we leveraged our mTORi-mediated in vitro developmental pausing system to achieve a time-resolved understanding of dormancy entry. By performing time-series proteomics analysis and modeling of the ESCs and trophoblast stem cells (TSCs) response to mTORi, we identify distinct pathways that are critical for the establishment of a reversible paused state. We find that the metabolic switch to fatty acid oxidation (FAO) not only enhances the maintenance of embryos in pause, but also shapes the epiblast morphology. By supplementing embryo culture with the FAO bottleneck metabolite L-carnitine, embryo longevity is extended up to 34 days in culture. We propose that facilitating the use of lipid reserves via L-carnitine supplementation enhances developmental pausing by establishing a deeper dormant state.

## Results

### Reversible pausing of ESCs, but not TSCs, in vitro

To investigate the tissue-specific restrictions to mTORi-induced pausing, we cultured ESCs and TSCs, which are the stabilized in vitro derivatives of the pre-implantation epiblast and trophectoderm, in mTORi^17-19^. As expected, ESCs significantly reduced proliferation and established a near-dormant state that retained colony morphology and marker expression of naive mouse ESCs (Fig. S1A)^13^. Importantly, ESCs reactivated upon withdrawal of mTORi without compromising stem cell colony morphology (Fig.S1A)^13^. In contrast, TSC colony morphology deteriorated over time and spontaneous differentiation was observed under prolonged mTORi and upon reactivation, despite the initial expected reduction in proliferation (Fig. S1A-B). Thus, ESCs and TSCs showed intrinsically different capacities to respond to mTOR inhibition and to maintain stemness under prolonged mTORi.

To isolate ESC-specific pathways that may allow their successful entry into pause, we performed a quantitative, time-resolved analysis of global ESC and TSC proteomes at the onset of mTORi-mediated entry and exit from pausing (denoted pause and release from here onwards respectively, Fig. 1A). Via label-free quantitative mass spectrometry of three biological replicates, we detected 7795 proteins (Data S1). Principal component analysis separated the samples primarily by cell type as expected (PC1, 38.8%, Fig. 1B). TSC pause and release samples further separated into two different clusters, while ESCs did not (PC2, 5.5%, Fig. 1B). This suggests that the TSC proteome underwent irreversible changes upon mTORi treatment, which is in line with the morphological changes (Fig. S1A). To further investigate proteome similarity at the molecular level, we constructed a pseudotime trajectory using the time-series proteome datasets. The pseudotime analysis also revealed distinct trajectories of ESCs and TSCs during pause and release (Fig. 1C, D). For ESCs, pseudotime and biological time largely correlated during pause and release (Fig. 1C), indicating a high degree of proteome similarity at distinct time points. Furthermore, pause and release trajectories anti-correlated and crossed paths, indicating reversibility of molecular changes. These data suggest a model where ESCs implement a defined sequence of events to establish pausing, and upon release from mTORi, revert these changes in the opposite order of their establishment, i.e., in a mirror-image fashion. As the release trajectory did not fully reach the starting point, a complete reversal likely takes more than 48 hours (Fig. 1C). Interestingly, the pause-release samples showed less variability than their parental ESCs grown in serum/LIF, suggesting that paused pluripotent cells may have more uniform pluripotency characteristics (Fig. 1C). On the contrary, TSCs did not show a clear pseudotime or reversible molecular behavior (Fig. 1D). Thus, not all stem cells undergo reversible dormancy in response to mTORi and this may be due to the inability to initiate a non-stochastic dormancy program.

**Figure 1.**
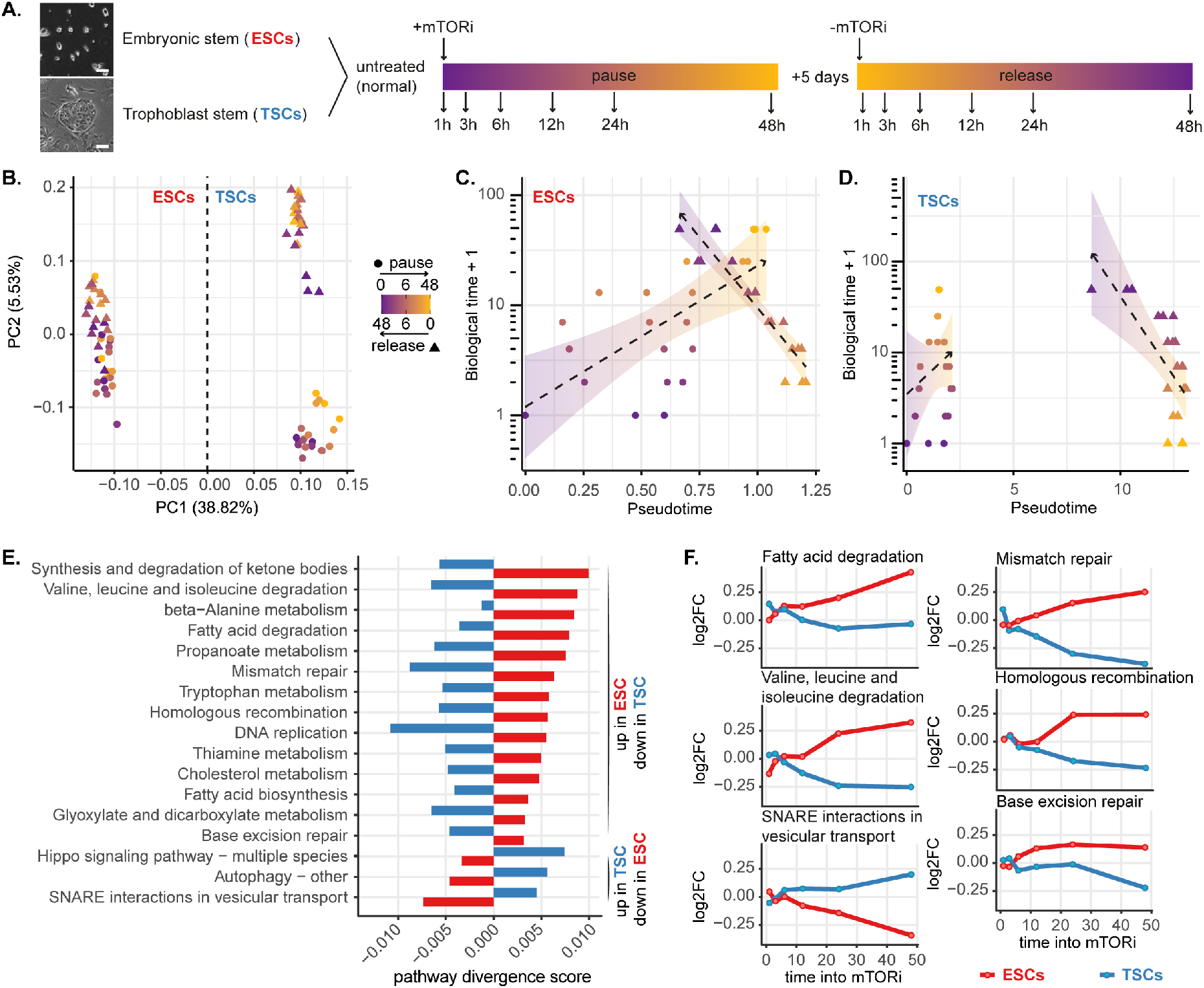
Pausing is reversible in ESCs, but not in TSCs. **(A)** Experimental setup: ESCs and TSCs were paused via mTORi for a total of 7 days, then released. Three biological replicates were collected at the indicated time points for proteomics analysis. Scale bar: 100 µm. **(B)** Principal component analysis of ESC and TSC proteomics data. The color scale indicates the time point during entry into pausing or release from pausing. **(C, D)** Pseudotime analysis of ESCs (C) or TSCs (D) indicates reversibility of pausing in the ESC proteome. **(E)** Divergent KEGG pathways during entry into pausing in ESCs and TSCs. The pathway score was computed based on mean expression of pathway proteins over time (see Materials and Methods). Selected pathways with a divergence score of > 0.0075 are displayed (see Data S2 for the full list). **(F)** Line plots of log_2_ fold change (FC) of mean protein expression (mTORi/control t=0) of selected divergent

### Pathway divergence in embryonic and extraembryonic cells in response to mTOR inhibition

Since ESCs reversibly paused and TSCs did not; we reasoned that pathways essential for reversible pausing can be isolated by comparing pathway usage in ESCs and TSCs over time. Direct comparison of each pathway per time point in ESCs vs TSCs revealed a widely heterogeneous distribution of pathway expression scores at each time point, thus showing a mixture of commonly used and divergent pathways (pathway expression score = mean protein expression in each KEGG pathway, Fig. S1C). To better distinguish between common and divergent pathways, we computed a pathway divergence score by following pathway expression over time in each cell type (Fig. 1E, F, S1D; see Methods for pathway divergence score).

A divergence score cut-off at 0.0075 yielded 25 divergent pathways in ESCs and TSCs during entry into pausing (Fig. 1E, F, Data S2). Metabolic pathways dominated the divergent behavior, among which, lipid and amino acid degradation were enriched (Fig. 1E). Interestingly, DNA repair pathways appeared to be enhanced in ESCs, but not in TSCs during pausing (Fig. 1F). Since several other metabolic pathways showed common patterns in ESCs and TSCs (Fig. S1E), these results suggest that metabolism of lipids and certain amino acids that show divergent patterns could be of specific relevance to establish developmental pausing.

### Immediate and adaptive changes en route to developmental pausing

To temporally resolve the steps in the ESCs’ entry into pause, we performed a time-series analysis of individual protein expression, followed by k-means clustering to identify temporal patterns (Fig. 2A, S2A for TSCs). We categorized the clusters as immediate or adaptive based on the expression patterns over time and performed gene ontology (GO) analysis on these categories (Data S3). Upon mTORi treatment, ESCs appeared to immediately reduce translation and ribosome biogenesis and reorganize chromatin (Fig. 2B-C). Interestingly, regulation of cell cycle, lipid metabolism, and DNA repair were among adaptive responses and may be enabled by prior steps in the establishment of pausing (Fig. 2B). TSCs did not upregulate chromatin organization-, DNA repair- or lipid metabolism-related proteins despite downregulating translation and ribosome biogenesis (Fig. 2D). These data suggest that the immediate response to mTORi, reduction of translation, is commonly realized in both cell types, yet TSCs lack the adaptive response to establish pausing.

**Figure 2.**
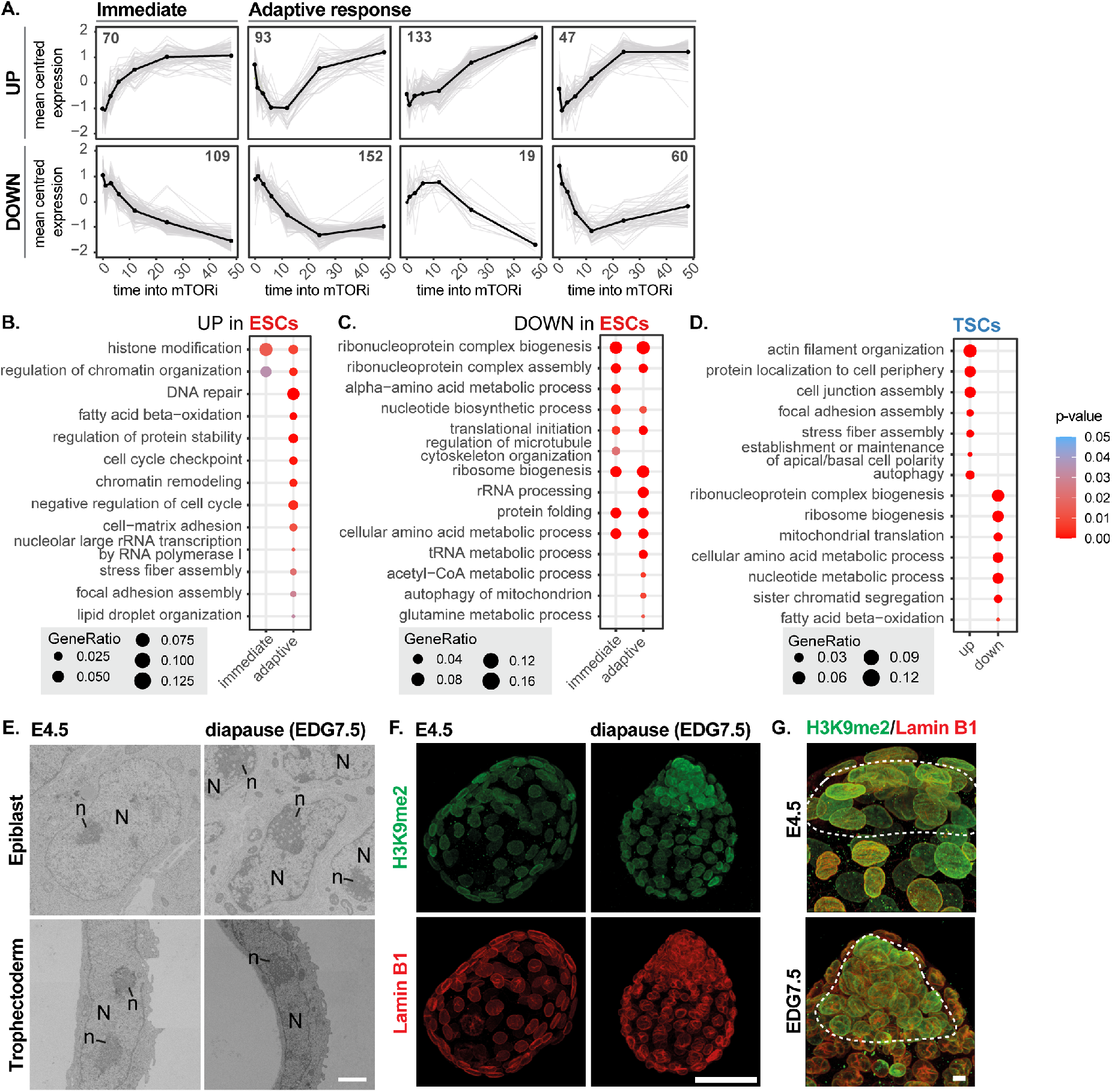
Orchestrated proteome rewiring during entry into pausing in ESCs. **(A)** Immediate and adaptive protein expression changes in ESCs. K-means clustering of differentially expressed proteins identified with MetaboAnalyst 4.0. **(B, C, D)** Gene ontology analysis of upregulated (B) and downregulated (C) proteins indicates that the immediately affected pathways histone modification and regulation of chromatin organization are also used during maintenance of pausing. Gene ontology analysis of proteome changes in TSCs (D). The full list of significantly enriched GO terms is provided in Data S3. **(E)** Transmission electron microscopy images of selected nuclei in the epiblast and TE of E4.5 and diapaused embryos. N = nucleus, n = nucleolus). Scale bar: 2 µm. **(F, G)** IF staining of H3K9me2 and Lamin B in E4.5 and diapaused (EDG8.5) embryos. Projections at 20x, in which the scale bar is 50 µm (F), and close-ups at 63x (G), in which the ICM is indicated in the dashed line.

To investigate whether changes at the protein level predictive of differential chromatin and adhesion properties of paused cells are reflected at the structural level, we performed transmission electron microscopy of whole blastocysts at E4.5 (n=4) or four days into diapause (n=4, at equivalent day of gestation (EDG) 7.5). Intriguingly, pluripotent epiblast cells showed extensive chromatin reorganization in diapause, with accumulation of heterochromatin at the nuclear periphery and denser, punctuated nucleoli, which are not seen in TE cells (Fig. 2E). The heterochromatin mark H3K9me2 was differentially enriched in the epiblast compared to TE, highlighting the tissue-specific chromatin reorganization (Fig. 2F-G). We also observed stress fibers and large focal adhesion complexes in diapaused embryos, as predicted by the proteome analysis (Fig. S2B). Thus, unlike in TSCs/ TE, establishment of pausing in ESCs and the epiblast appear to involve immediate rewiring of the chromatin landscape and reduced protein synthesis, followed by a metabolic switch to the usage of lipids.

### Enhancing lipid usage via carnitine supplementation boosts survival of in vitro paused embryos

Our proteomic analysis points to metabolism, particularly the degradation of lipids and certain amino acids, as a means to establish a paused state in ESCs (Fig. 1E-F, 2B). Real-time readout of global metabolic activity via Seahorse analysis showed a precipitous decrease in the rate of glycolysis, as well as basal respiration in paused ESCs, while TSCs displayed a milder decrease (Fig. S3A). Pausing may necessitate metabolic rewiring in ESCs, but not in TSCs, since TSCs have lower metabolic activity compared to ESCs even before mTORi treatment. To more precisely pinpoint metabolic alterations and identify critical metabolites in paused ESCs, we next performed bulk metabolomics (Fig. 3A, Data S4). We found that the differentially enriched metabolites were highly enriched for lipid molecules, corroborating the proteome analysis. Specifically, paused ESCs accumulated long chain carnitine-conjugated fatty acids, ceramide, and sphingomyelins; but were depleted of short-chain carnitine-conjugated fatty acids (Fig. 3A). FAO provides energy but not many building blocks for proliferation and thus may be an ideal energy pathway in dormant cells. Proteins involved in FAO were upregulated ∼24 hours after mTORi treatment and were downregulated upon release from pausing, while glycolysis genes did not show this dynamic pattern (Fig. 3B, only significantly changed proteins are shown). Thus, FAO is likely the main energy pathway in paused pluripotent cells.

**Figure 3.**
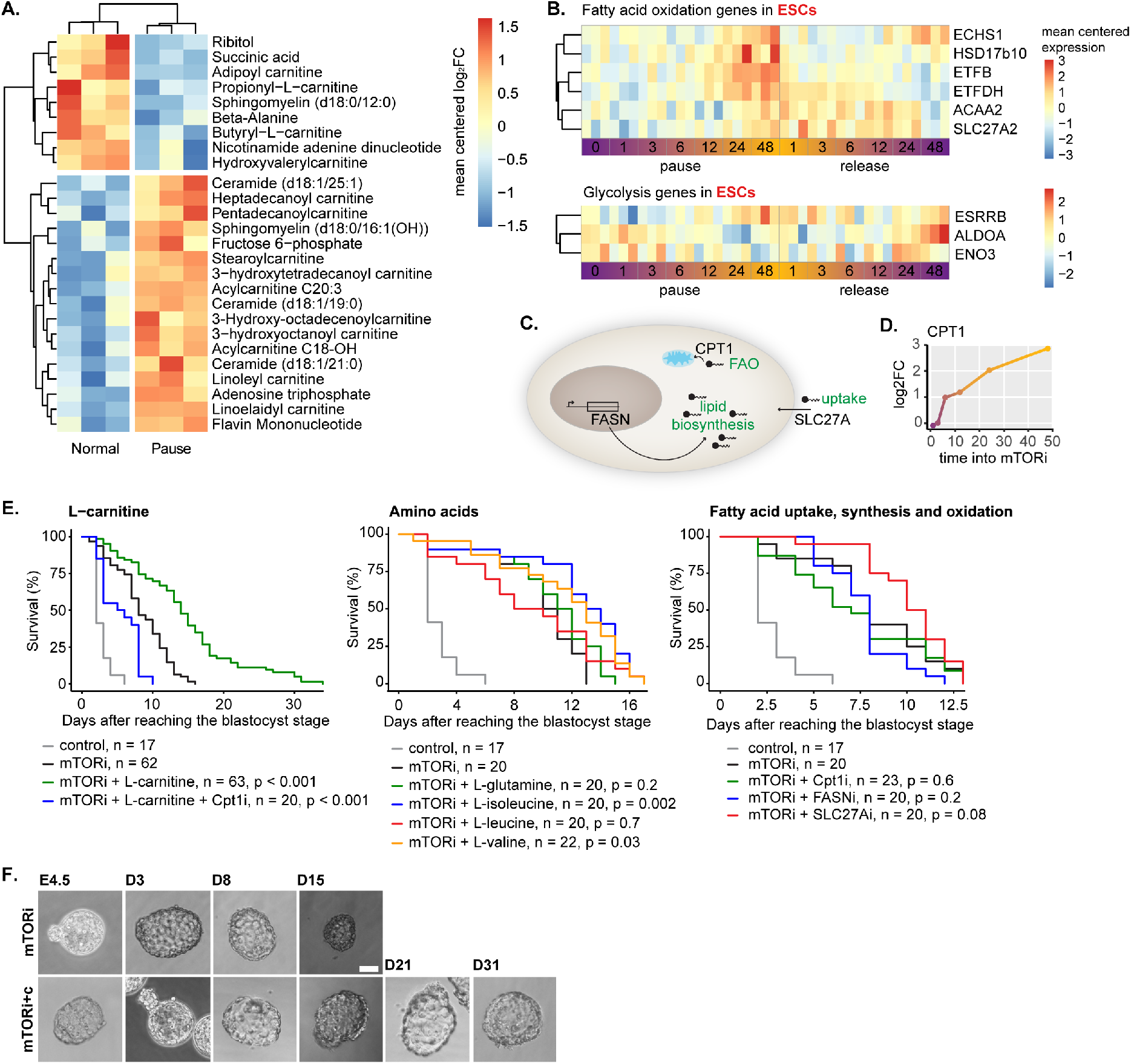
L-carnitine supplementation significantly enhances in vitro developmental pausing. **(A)** Differentially expressed metabolites (p-value <0.05 and absolute log2FC >0.75) in paused (7 days) vs normal ESCs. Mean centered data of three biological replicates are shown. **(B)** Expression levels of differentially expressed fatty acid oxidation and glycolysis-related genes in ESCs. Mean centered protein expression of three biological replicates are shown. **(C)** Schematic overview of fatty acid uptake, synthesis, and oxidation, in which processes are indicated in green and respective genes in black. **(D)** Cpt1a protein expression into mTORi in ESCs. **(E)** Embryo survival curves in the shown conditions. n = number of embryos. Statistical test is the G-rho family test of of Harrington and Fleming^59^. **(F)** Bright field images of mTORi and carnitine supplemented paused embryos. Scale bar: 50 µm.

Transfer of fatty acids from the cytosol into mitochondria requires their conjugation to carnitine by CPT1 (Fig. 3C)^20^. This step is tightly regulated and acts as a bottleneck in FAO capacity. CPT1 expression increases upon ESC pausing (Fig. 3D). We reasoned that the survival of in vitro-paused blastocysts may be restricted by the depletion of free carnitine (Fig. 3A), leading to inefficient usage of lipids. In agreement with this idea, supplementation of blastocysts with free L-carnitine during mTORi-mediated pausing remarkably enhanced their pausing duration (Fig. 3E-F, mTORi-only median and maximum survival = 8 and 15 days, mTORi+carnitine median and maximum survival = 15 and 34 days). Inhibition of CPT1^21^ canceled the beneficial effect of L-carnitine supplementation, indicating that the benefit is through FAO (Fig. 3E). Direct supplementation of carnitine-coupled short-chain fatty acids and free fatty acids in culture media was mostly toxic to embryos and did not extend pausing beyond mTORi treatment, except for adipoyl-L-carnitine (Fig. S3B). As short chain carnitines can be generated from branched-chain amino acids, and this amino acid degradation pathway is specifically enriched in paused ESCs (Fig. 1F)^22^, we also tested whether amino acid supplementation enhances pausing (Fig. 3E). Isoleucine and valine significantly enhanced pausing, although not to the extent of direct L-carnitine supplementation. Inhibition of fatty acid uptake via SLC27Ai^23^ or synthesis via FASNi^24^ did not alter maximum survival, suggesting that stored cellular lipids are used in paused cells (Fig. 3C, E). Supplementation with the citric acid cycle intermediate ɑ-ketoglutarate or vitamin C as antioxidant also did not significantly enhance pausing, ruling out the contribution of other factors downstream of fatty acid oxidation (Fig. S3B). Taken together, L-carnitine supplementation strikingly enhanced developmental pausing in vitro through fatty acid oxidation.

### Balanced lipid usage in carnitine-supplemented embryos

Proteomic analysis suggested the tissue-specific utilization of lipids in paused cells. We next investigated lipid usage in in vivo- and in vitro-paused embryos via electron microscopy and fluorescence staining of lipid droplets (LD). Electron microscopy revealed a striking difference in lipid droplet usage in the epiblast and TE of in vivo diapaused embryos (Fig. 4A). While the epiblast and TE had several small LDs in E4.5 embryos, each TE cell of diapaused embryos in most cases had a single, very large LD (Fig. 4A). On the contrary, the epiblast LDs remained in a similar size range (Fig. 4A). Through manual annotations of LDs and mitochondria in single sections from four separate blastocysts in each condition, we found that the overall number of LDs decreased in diapause, while their average size increased (Fig. 4B, S4A-C). More mitochondria were located in the vicinity of LDs, pointing to their active usage (Fig. 4B). Fluorescence imaging of LDs through Bodipy staining confirmed the presence of large LDs in TE, but not the epiblast in diapause (Fig. 4C). Interestingly, large LDs were found on the mural side, but not the polar side of the blastocyst, indicating a heterogeneous and imbalanced metabolic pattern in the TE.

**Figure 4.**
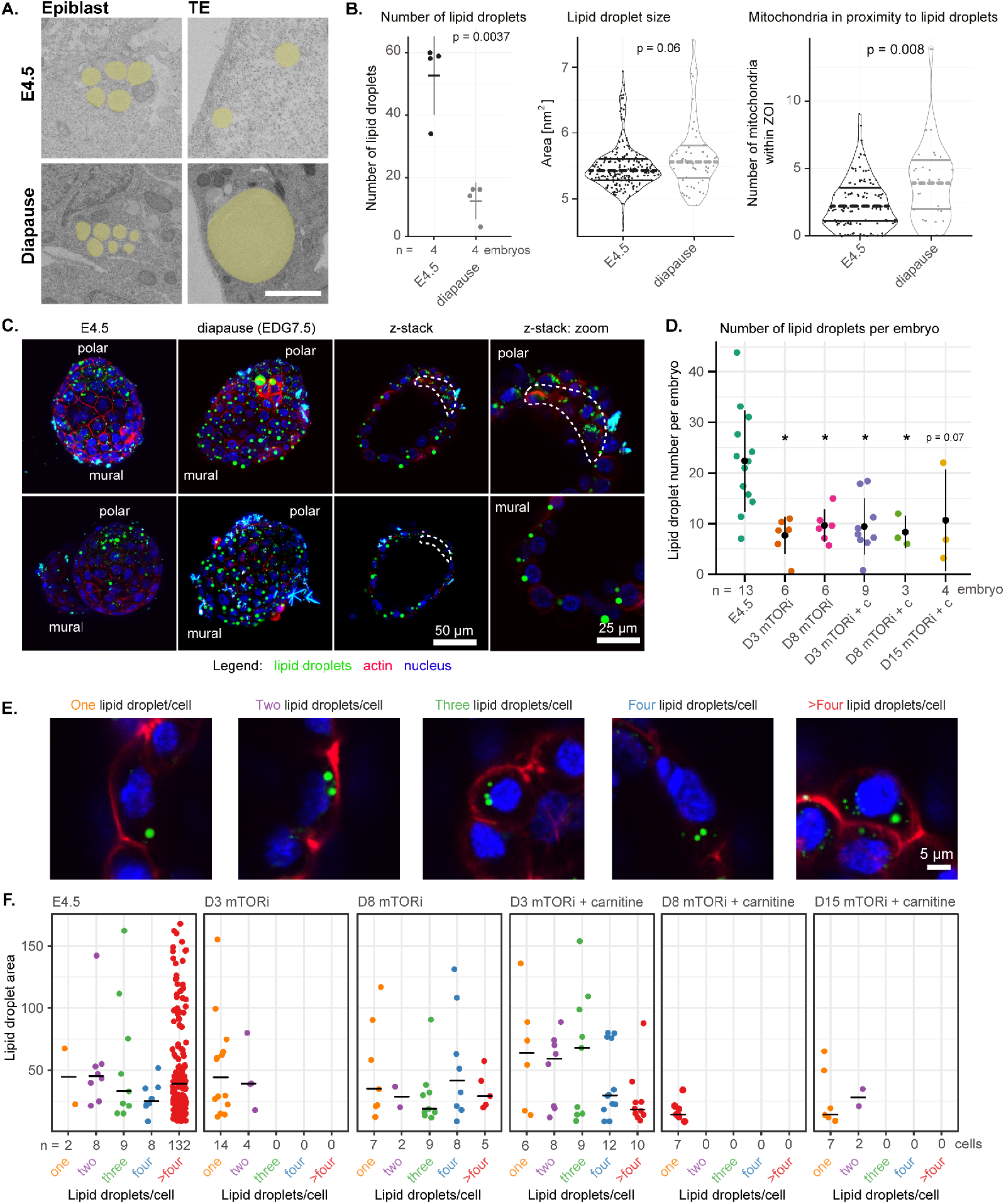
L-carnitine facilitates lipid usage during developmental pausing. **(A)** Electron microscopy (EM) images of E4.5 and diapaused embryos. Lipid droplets are highlighted in yellow. Scale bar: **2** µm. **(B)** Quantifications of LD abundance and size and the number of mitochondria in proximity to LDs. Whole-blastocyst EM images (single plane) from four individual E4.5 or diapaused embryos were used. **(C)** Visualization of LDs in E4.5 or diapaused embryos via Bodipy staining (green). Red channel shows an actin stain (CellMask). **(D)** Number of LDs per embryo in the shown conditions. Live embryos were stained with Bodipy (LDs), CellMask (actin) and Hoechst (DNA). n = number of embryos per condition. *: p <0.05, one way ANOVA, Dunnett’s post hoc test. **(E)** Representative images of cells with one, two, three, four, or >4 LDs. LDs are shown in green, cell

We next investigated LD abundance and size in carnitine-supplemented embryos via Bodipy staining, followed by image-based quantification of LDs per cell (Fig. 4D-F). As fixing the embryos impaired the Bodipy staining, a tissue-specific LD quantification was not possible. In general, in vitro-paused embryos with or without carnitine had fewer LDs similar to in vivo diapaused embryos (Fig. 4D). However, carnitine altered the pattern of lipid droplet abundance and size, indicating altered lipid usage. mTORi-only embryos seem to switch to using lipids after day 3, as seen via the dispersion of lipids into multiple LDs (Fig. 4F, D3 vs D8 mTORi panels; representative images of one, two, three, four, and >4 lipid droplets per cells are shown in Fig. 4E). Carnitine-supplemented embryos, however, showed this pattern already on day 3, suggesting a balanced lipid usage already in the early phases of pausing. At day 8 and day 15, the vast majority of cells contained one-two small LDs. (Fig. 4F). By day 15, the majority of mTORi-only embryos collapsed, while 50% of carnitine-supplemented ones were still alive (Fig. 3E). Collectively, these data suggest that L-carnitine prolongs in vitro developmental pausing by balancing lipid usage.

### Reduced genomic and metabolic activity of carnitine-supplemented embryos

We have previously shown that mTORi-paused embryos reside in a near-dormant state with low nascent transcription, histone acetylation and translation^13^. Yet, the TE showed higher genomic activity than the inner cell mass (ICM)^13^. Fatty acid degradation directly contributes to the cellular quantity of acetyl-CoA, thereby providing acetyl donors for histone acetylation^25^. Quiescent cells have been associated with low or high histone acetylation in different contexts, and the histone acetyltransferase MOF has been directly implicated in regulating quiescence^26^. To address whether lipid usage influences global histone acetylation levels in the TE and ICM of carnitine-supplemented embryos, we stained the embryos for histone H4 lysine 16 acetylation (H4K16ac) and performed image-based quantification (Fig. 5).

**Figure 5.**
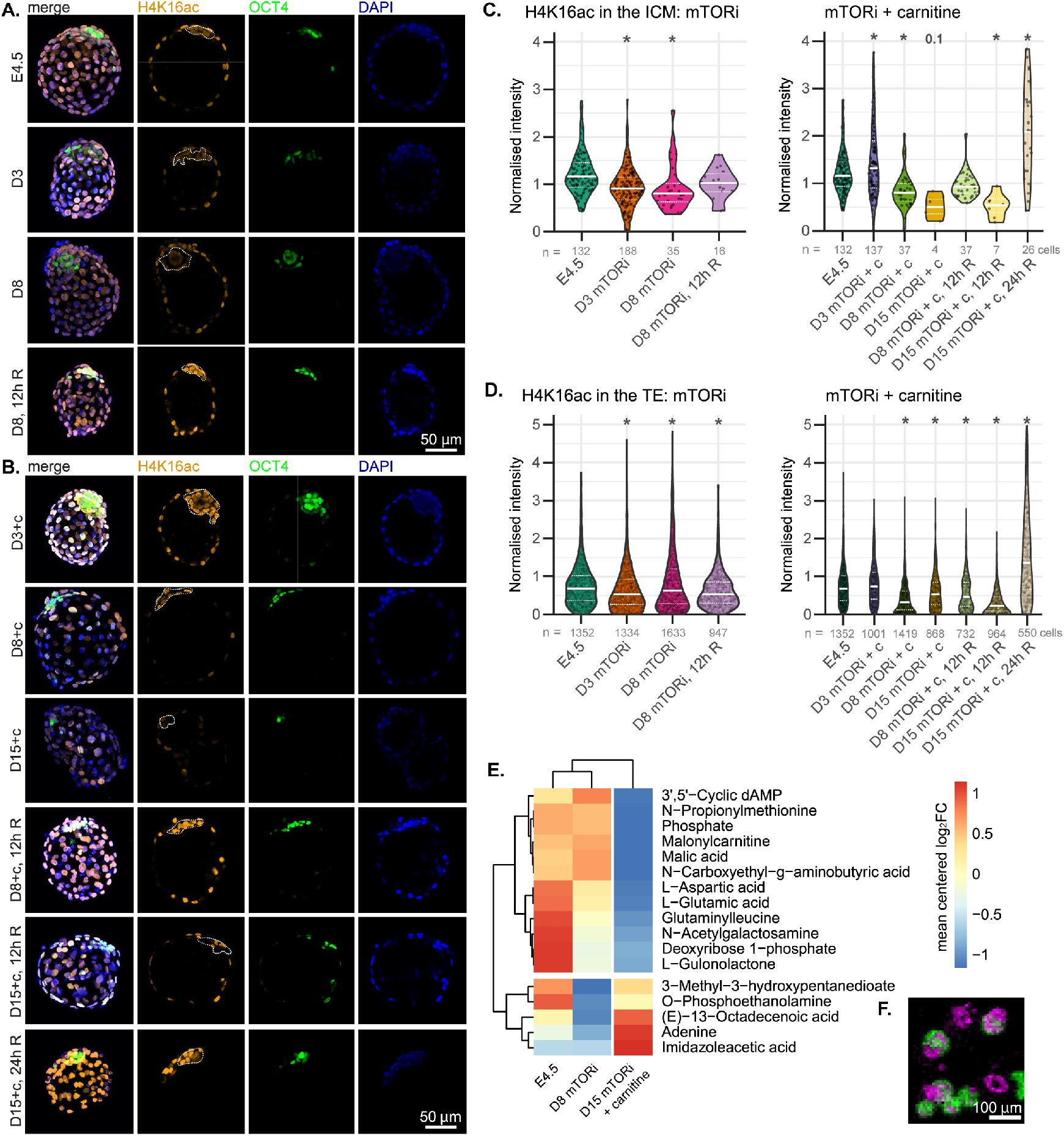
Carnitine supplementation reduces global genomic and metabolic activity. **(A, B)** IF staining of H4K16ac and OCT4 in E4.5, mTORi (A), and mTORi+carnitine (mTORi+c) embryos (B). Embryos (n=8-12/condition) were paused for 3, 8 or 15 days and or reactivated (R) for 12 hours or 24 hours in fresh medium without mTORi. For each condition, a representative embryo’s z-projection and single z-stack channels are shown. The ICM is highlighted with a dashed white line. Scale bar = 50 µm. **(C, D)** Quantification of DAPI-normalized H4K16ac intensities in the ICM (OCT4-positive cells, C) or TE (D) in each condition. n = number of cells, *: p <0.05, one way ANOVA, Dunnett’s post hoc test. **(E)** Mean centered log_2_FC of metabolites detected in single embryo MALDI-imaging metabolomics analysis (n = 4-12 embryos/condition/metabolite). **(F)** An example image of the single embryo MALDI-imaging, showing two metabolites in green and purple. Scale bar = 100 µm.

Paused embryos with or without carnitine showed a global decrease in H4K16ac levels, except for a transient increase in D3 mTORi+carnitine embryos (Fig. 5A-D). The ICM showed a more dynamic response compared to TE, with gradual reduction in H4K16ac until D8 of mTORi and D15 of mTORi+carnitine embryos (Fig. 5C-D). Overall, mTORi+carnitine embryos showed lower H4K16ac at later time points compared to mTORi embryos. Thus, carnitine supplementation seemed to facilitate a deeper dormant state. Moreover, proliferation ceased from the mural to polar side of the embryos in both mTORi alone and carnitine supplemented embryos (Fig. S5D-E). Single embryo metabolomics through MALDI-imaging mass spectrometry also showed globally reduced metabolic activity in mTORi+carnitine embryos compared to E4.5 and even mTORi embryos (Fig. 5E, F). Although the number of metabolites that are detectable at the single embryo level is limited, there was little variation between individual embryos for detected metabolites (Fig. S5F). Thus, carnitine-supplemented mTORi-paused embryos seemed to be genomically and metabolically dormant.

To test whether mTORi+carnitine embryos were still able to reactivate and subsequently increase global genomic activity, we withdrew mTORi and carnitine from embryo culture and allowed the embryos to reactivate for 12 or 24 hours (denoted as + 12/24h R). Both D8 mTORi and D8 mTORi+carnitine embryos were able to reactivate and showed high H4K16ac levels after 12 hours (Fig. 5A-D). D15 mTORi+carnitine embryos showed only initial signs of reactivation after 12 hours and full reactivation after 24 hours (Fig. 5B-D). Overall, there was a reduction in OCT4-positive cells, as well as TE cells in both mTORi and mTORi+carnitine embryos (Fig. S5A, B), however we did not observe high levels of apoptosis (Fig. S5C). Levels of OCT4 and NANOG are regulated by the translational capacity of pluripotent cells, therefore the global reduction of translation as a result of mTOR inhibition reduces their levels^27^. This was however reversible upon reactivation, as the number of OCT4-positive cells increased after reactivation of embryos (Fig. 5B, S5A). Intriguingly, we noticed rosette-like structures in the ICM of both mTORi and mTORi+carnitine embryos (Fig. 5B, D3+C embryo). The epiblast of in vivo diapaused embryos was organized into a rosette-like structure around EDG9.5 (equivalent to D6 mTORi in this study), following a preceding increase in WNT pathway activity^28^. Here we found that 12 and 20% of mTORi embryos show a rosette-like structure on days 3 and 8, respectively (one out of eight on D3 and two out of ten on D8, Fig. 5A). mTORi+carnitine embryos generated rosette-like structures more efficiently and earlier than mTORi only (four out of eight (50%) on D3 and zero out of eleven (0%) on day 8, Figure 5B). Whether the ICM retained the rosette-like structure after D3 is unclear due to the reduced OCT4 staining. These results point to a direct link between lipid metabolism and epiblast morphology and suggest that carnitine-supplementation may better recapitulate the ICM morphology of in vivo embryos.

### FOXO1 activity is essential for carnitine enhanced developmental pausing

Forkhead box (FOX) family transcription factors (known as DAF16 in C.elegans) regulate cellular metabolism, nutrient stress, longevity and lifespan in multiple species^29-31^ and show dynamic cellular localization in response to cellular signaling^32,33^. In addition, FOXO proteins have been proposed to play a role in marsupial and killifish diapause^34,35^.

As FOXO1 is a known regulator of lipid metabolism^36,37^, and was upregulated in mTORi-treated ESCs and TSCs (Fig. S6A), we reasoned that it might be a central player in the metabolic rewiring during developmental pausing. Chemical inhibition of FOXO1^38^ interfered with mTORi-mediated pausing in a dose-dependent manner and completely abolished pausing at 1 µM, with or without carnitine supplementation (Fig. 6A, S6B, used concentrations are not cytotoxic). Therefore, enhancement of pausing by L-carnitine was dependent on FOXO1 activity. Immuno-fluorescence staining revealed that carnitine supplementation enhanced FOXO1 expression in the ICM (Fig. 6B-D). Surprisingly, the rosette-like epiblast showed cytoplasmic FOXO1 localization despite mTOR inhibition. Thus, FOXO1 may have distinct roles at different stages of pluripotency and pausing. Taken together, we show the critical role of FOXO1-mediated lipid metabolism in the maintenance of cellular dormancy during developmental pausing and reveal intriguing links between lipid metabolism, pluripotency status and embryonic morphology (Fig. 6F).

**Figure 6.**
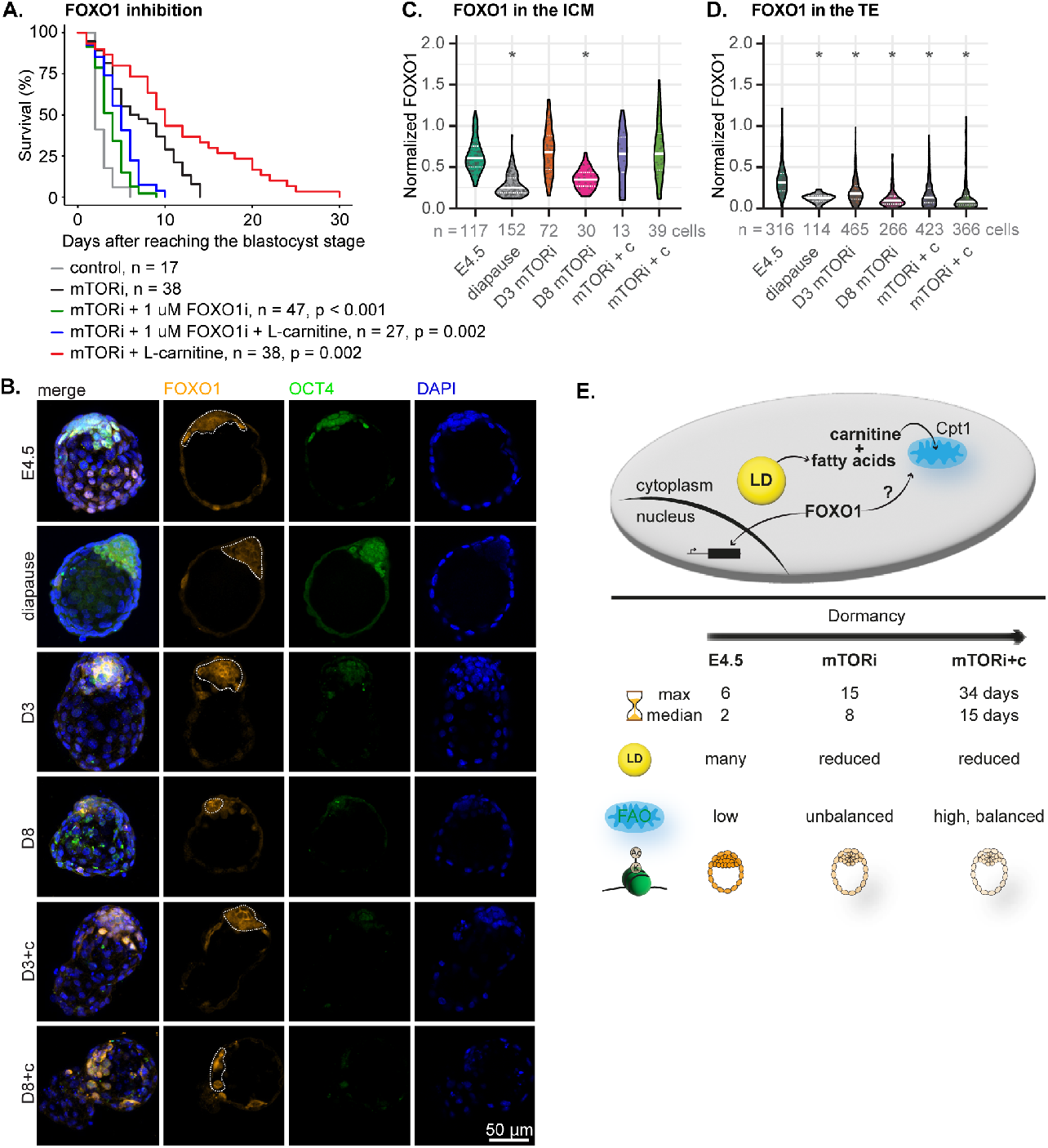
FOXO1 activity is essential for carnitine-enhanced developmental pausing. **(A)** Embryo survival curves in the shown conditions. FOXO1 inhibition completely reverses carnitine-enhanced developmental pausing. n = number of embryos. **(B)** IF staining for FOXO1 and OCT4 in each condition. Embryos (n=7-8/condition) were paused for 3 or 8 days with or without L-carnitine. Diapaused embryos were retrieved at EDG8.5 (n=4). Representative embryos’ z-projection and single z-stack of each channel is shown. The ICM is highlighted with a dashed white line. Scale bar = 50 µm. **(C, D)** Quantification of area normalized FOXO1 intensity in the ICM (C) or TE (D) in the shown conditions. n = number of cells, *: p <0.05, one way ANOVA, Dunnett’s post hoc test.

## Discussion

In this study, we investigated the cell type-specific response to mTORi-induced in vitro developmental pausing. By comparing pathway expression in ESCs and TSCs, we identified lipid metabolism as an important regulator of dormancy, embryo longevity, as well as epiblast morphology. The reorganization of epiblast cells into a rosette-like structure (at ∼EDG9.5, D6 of diapause) followed a transient increase in Wnt signaling (at ∼EDG7.5) in diapaused embryos^28^. In vitro paused embryos show epiblast polarization similar to in vivo diapaused embryos, although large variations in embryo shape and morphology are seen. A subset of mTORi-paused embryos showed rosette-like morphology on D8 (this study), while carnitine-supplemented ones showed it already on D3 of pausing. The earlier onset of polarization in carnitine supplemented embryos at D3 points to a direct link between fatty acid oxidation and Wnt pathway activity. WNT proteins are lipid-modified to enable their secretion^39^. Interestingly, inhibition of WNT secretion, and thus paracrine signaling, reduces Wnt pathway activity without compromising polarization capacity, but alters its timing (EDG7.5 instead of 9.5)^28^. Carnitine supplementation may thus influence the timing of epiblast polarization by modulating WNT secretion.

Lipid consumption has been associated with a naive-to-primed pluripotency transition^40,41^. The early onset of polarization in carnitine-supplemented embryos supports this notion. The diapaused epiblast is thought to reside in a naive pluripotent state, however only EDG6.5 embryos were used for the transcriptomics profiling underlying this conclusion^14^. Time - series transcriptomics analysis of in vivo- and in vitro-paused embryos is likely to reveal dynamic progression of pluripotency during diapause. The mouse epiblast normally polarizes only after implantation (∼E5.0-5.25)^42^. The early-onset polarization of in vivo- and in vitro-paused epiblast, and particularly in mTORi+carnitine, indicates that neither implantation, nor a specific stage of TE differentiation are prerequisites to polarization. Whether polarization is required for epiblast maintenance in prolonged diapause is worth investigating in further studies.

Lipid reserves and FAO appear to play an essential role in regulating the balance of stem cell dormancy and proliferation in several adult tissues^8,43-46^, as well as during embryonic diapause across species^35,47-49^, although a contrary observation has been made in human cells^26^. Interestingly, enhanced lipid consumption due to dietary restriction increases life span in Drosophila ^50^ and C.elegans^51^. Similarly, one of the most prominent regulators of lipid metabolism, FOXO1 (DAF16 in C.elegans), is associated with longevity^31^. Involvement of mTOR, FOXO1 and FAO in both diapause and lifespan regulation points to a partially shared regulatory network between these two phenomena. Thus, further exploration of diapause biology will likely reveal regulators of longevity.

A further prolongation of developmental pausing would require lipid supplementation, as diapaused mouse embryos eventually deplete lipid reserves^48^ and do not seem to de novo synthesize sufficient lipids to fulfil the energy requirement^48,52^. However, our data indicate that other lipids than short chain fatty acids should be supplemented during developmental pausing. Alternative delivery methods including delivery of long chain triglycerides or solid lipid nanoparticles^53^ would offer opportunities to replenish the fatty acid reserves, thereby extending the current maximum survival duration of 34 days. Replenishing lipid reserves would not only further enhance developmental pausing, but may also affect membrane fluidity through altering the fatty acid composition^54-56^. As membrane fluidity of cells or the extracellular matrix influences stem cell maintenance and differentiation ^57,58^, the relationship between FAO, altered membrane fluidity, and maintenance of stemness warrants further exploration.

Although dormancy as an end state is simply characterized by loss of proliferation ^1^, specific steps taken to establish dormancy remained unclear. Here we show that the immediate next step of mTOR inhibition, i.e., reduction in protein synthesis, does not suffice to establish a dormant state because TSCs reduce translational proteins, however, do not successfully establish reversible dormancy. We propose that active chromatin rewiring, followed by a metabolic shift to FAO are critical steps in the establishment of dormancy. Increased heterochromatin formation either at the nuclear lamina or around nucleoli may facilitate selective and transient silencing of differentiation and proliferation genes.

Together, our findings provide novel insights into the metabolic regulation of cellular dormancy during developmental pausing. We anticipate that studying how metabolism regulates pluripotency and cellular dormancy will provide a novel framework for understanding stem cell quiescence and longevity, with future implications for reproductive strategies of endangered species.

## Acknowledgments

We thank members of the Bulut-Karslioglu Lab and Berna Sözen for discussions and feedback; MPIMG transgenics facility and animal house units for excellent technical support and discussions. We thank René Buschow for help with EM image analysis, Jennifer Shay and Cordula Mancini for assistance, Beata Lukaszewska-McGreal for proteome sample preparation, Rainer Glauben, Cansu Yerinde, and Werner Kammerloher for guidance in Seahorse assays.

## Funding

Swiss National Science Foundation Early Postdoc.Mobility fellowship P2EZP3_195682 (VAvdW)

Max Planck Society (ABK)

Sofia Kovalevskaja Award of the Humboldt Foundation (ABK)

ERC Consolidator grant METACELL, grant number 773089 (TA)

## Author contributions

ABK and VAvdW conceived and developed the project. DM performed proteomics and metabolomics runs. MS and BF performed electron microscopy sample preparation and imaging with input and supervision from TM. MS performed EM and embryo metabolomics image analysis. DPI performed Seahorse experiments. SR performed reversibility analysis. ABK collected proteomics and metabolomics samples. VAvdW performed all other experiments and analysis. ABK supervised the project. VAvdW, MS, and AB-K wrote the manuscript with feedback from all authors.

## Data and materials availability

Proteomics and metabolomics datasets generated in this study have been deposited to the PRIDE and peptideAtlas databases and will be made accessible upon publication.

## Materials and Methods

### Animal experimentation

All animal experiments were performed according to local animal welfare laws and approved by local authorities (covered by LaGeSo licenses ZH120, G0284/18, and G021/19). Mice were housed in individually ventilated cages and fed ad libitum.

### Cell lines and culture conditions

#### Mouse ESCs

E14 ESCs (received from S. Kinkley Lab, MPIMG) were used. Cells were plated on 0.1% gelatin-coated dishes and grown in DMEM high glucose with Glutamax media (Thermo, 31966047) supplemented with 15% FBS (Thermo, 2206648RP), 1x NEAA (Thermo, 11140-035), 1x β-mercaptoethanol

(Thermo, 21985023), 1x Penicillin/ streptomycin (Life Technologies, 15140148) and 1000U/mL LIF and grown at 37°C in 20% O2 and 5% CO2 incubator. At each passage, cells were dissociated using TrypLE (Thermo Fisher, 12604-021) with media change every day. Cells were routinely checked for mycoplasma.

#### Mouse TSCs

TSCs (received from M. Zernicka-Goetz Lab) were grown on mitotically-inactivated mouse embryonic fibroblasts in media containing RPMI1640+GlutaMAX (Gibco 61870010, Thermo Fisher), 20% FBS, 1x β-mercaptoethanol (Thermo, 21985023), 1x Penicillin/streptomycin (Life Technologies, 15140148), 1x sodium pyruvate, 25ng/µl FGF4 (R&D Systems #235-F4-025) and 1 µg/ml Heparin (Sigma cat # H3149). Cells were depleted off feeders at the time of collection for analysis.

### Developmental pausing setup

#### Mouse ESC/TSC pausing

For pausing of mouse ESCs and TSCs, cells were treated with the mTOR inhibitor INK-128 at 200 nM final concentration. Media was replenished daily.

#### In vitro fertilization

10- to 12-week-old b6d2F1 mice were superovulated by an intraperitional injection with PMSG (5IU/ 100 µl) on day 0 of the superovulation protocol. On day 2, mice were intraperitonially injection with HCG 5 IU/ 100 µl. On day 3, mice were sacrificed with CO2 and cervical dislocation, after which the cumulus oocyte complexes (COCs) were retrieved. A frozen sperm straw was removed from the liquid nitrogen and held in the air for 5 seconds, after which the straw was kept at 37°C for 10 minutes. Ten µl of sperm suspension was pipetted into the centre of 90 µl of Fertiup™ Mouse Sperm Preincubation Medium (CosmoBio, cat. # KYD-002-EX). Ten µl of motile sperm was added to each drop of fertilizing CARD Medium (CosmoBio, cat. # KYD-003-EX) containing the COCs. After overnight culture, the obtained 2-cell stage embryos were transferred to a fresh drop of KSOM (Merck, cat. # MR-107-D). The IVF with b6d2F1 oocytes and b6casF1 has on average a 2-cell development (fertilization) rate of 80%. From these around 70-75% develop to blastocysts.

#### Mouse blastocyst pausing in vitro

From developmental day 3.5, embryos were cultured in reduced KSOM medium made in-house (Table S1). 2x homemade reduced KSOM medium was prepared and stored for up to 3 months at -80°C. The medium was filtered through a 0.22 µm filter (Corning, cat. # 431118). 1x reduced KSOM medium was freshly prepared from the 2x frozen stock prior to each experiment and only after thawing, 100x CaCl2•2H2O and bovine serum albumin fraction V was added. The reduced KSOM contains a total nutrient content of 10.4 mM, compared to 13.5 mM in commercially available KSOM^60^. Embryos were cultured in 4-well dishes (Nunc IVF multidish, Thermo Scientific, cat. # 144444) in a volume of 500 µl reduced KSOM under hypoxic conditions (5% O2 and 5% CO2 at 37°C). Embryo survival was evaluated daily under a stereomicroscopy and dead (collapsed) embryos were removed.

#### Embryo supplements

Embryonic developmental day 3.5 embryos were cultured in reduced KSOM supplemented with single or combinations of supplements (Table S2). The supplement solvent concentration did not exceed 2%.

#### Survival analysis

Embryo survival curves were plotted in RStudio (version 1.3.1093 with R version 3.6.3) with the survminer (version 0.4.9) package. The survival (version 3.3-1) package was used to extract median and maximum survival values.

#### In vivo diapause induction

In vivo diapause was induced as previously described^61^ after natural mating of CD1 mice. Briefly, pregnant females were ovariectomized at E3.5 and afterwards injected every other day with 3 mg medroxyprogesterone 17-acetate (subcutaneously). Diapaused blastocysts were flushed from uteri in M2 media after 3 and 4 days of diapause at EDG7.5 and EDG8.5, respectively.

### Global proteomics

#### Sample preparation

Proteomics sample preparation was done according to a published protocol with minor modifications^62^. In brief, 5 million cells in biological duplicates were lysed under denaturing conditions in 500 µl of a buffer containing 3 M guanidinium chloride (GdmCl), 10 mM tris(2-carboxyethyl) phosphine, 40 mM chloroacetamide, and 100 mM Tris-HCl pH 8.5. Lysates were denatured at 95°C for 10 min shaking at 1000 rpm in a thermal shaker and sonicated in a water bath for 10 min. 100 µl lysate was diluted with a dilution buffer containing 10% acetonitrile and 25 mM Tris-HCl, pH 8.0, to reach a 1 M GdmCl concentration. Then, proteins were digested with LysC (Roche, Basel, Switzerland; enzyme to protein ratio 1:50, MS-grade) shaking at 700 rpm at 37°C for 2 hours. The digestion mixture was diluted again with the same dilution buffer to reach 0.5 M GdmCl, followed by a tryptic digestion (Roche, enzyme to protein ratio 1:50, MS-grade) and incubation at 37°C overnight in a thermal shaker at 700 rpm. Peptide desalting was performed according to the manufacturer’s instructions (Pierce C18 Tips, Thermo Scientific, Waltham, MA). Desalted peptides were reconstituted in 0.1% formic acid in water and further separated into four fractions by strong cation exchange chromatography (SCX, 3M Purification, Meriden, CT). Eluates were first dried in a SpeedVac, then dissolved in 5% acetonitrile and 2% formic acid in water, briefly vortexed, and sonicated in a water bath for 30 seconds prior to injection to nano-LC-MS/MS.

#### Run parameters

LC-MS/MS was carried out by nanoflow reverse phase liquid chromatography (Dionex Ultimate 3000, Thermo Scientific) coupled online to a Q-Exactive HF Orbitrap mass spectrometer (Thermo Scientific), as reported previously^63^. Briefly, the LC separation was performed using a PicoFrit analytical column (75 μm ID × 50 cm long, 15 µm Tip ID; New Objectives, Woburn, MA) in-house packed with 3-µm C18 resin (Reprosil-AQ Pur, Dr. Maisch, Ammerbuch, Germany).

#### Peptide analysis

Raw MS data were processed with MaxQuant software (v1.6.10.43) and searched against the mouse proteome database UniProtKB with 55,153 entries, released in August 2019. The MaxQuant processed output files can be found in Data S1, showing peptide and protein identification, accession numbers, % sequence coverage of the protein, q-values, and label free quantification (LFQ) intensities. The mass spectrometry data have been deposited to the ProteomeXchange Consortium (http://proteomecentral.proteomexchange.org) via the PRIDE partner repository^64^ with the dataset identifier PXD033750 and PXD033798.

#### Global proteomics analysis

The DEP package (version 1.14.0)^65^ was used for the global proteomics analysis. Potential contaminants were filtered, unique gene names were generated, and only proteins that were quantified in two out of three replicates were included for further analysis. Data was normalized and missing values were imputed using random draws from a Gaussian distribution centered around a minimal value. A total of 7795 proteins was detected and quantified in both ES and TSCs. Protein dynamics were plotted using a principal component analysis (PCA).

#### Pseudotimeanalysis

Unique peptide abundances were first averaged across replicates. Then, the 500 most variable proteins were selected as determined by their variance for each cell type and condition and used for downstream analysis. The z-score was calculated for each protein, cell type and condition. Diffusion maps were then calculated on the z-scores using the destiny package version 3.8.0 in R version 4.1.0 with parameter n_pcs = 10. From these diffusion maps we calculated a diffusion pseudotime using the DPT function of the destiny package.

### Pathway divergence analysis

Differentially expressed genes were identified using the DEP package. Data were prepared and filtered as described under the global proteomics analysis. Log2FC and adjusted p-values during entry into pausing for all proteins in both ES and TSCs compared to 0h were computed. KEGG^66^ pathways containing at least 10 genes symbols were included in the divergence analysis. The pathway expression value was defined as the mean log2FC of proteins between any of the time points during entry and the 0h control. A total of 327 pathways had a positive or negative mean log2FC pathway expression value in ESCs at any of the time points during entry compared to the 0h control (Data S2).

Divergence score was calculated as DS = mESC (mean log2FC pathway expression tP/tN)-mTSC (mean log2FC pathway expression tP/tN)

where, P = pause time points, N=normal (untreated), and m denotes the slope of pathway expression over time, computed with linear regression.

Pathways with a divergence score of > 0.0075 were considered divergent and plotted with a waterfall plot using ggplot2 (version 3.3.5). Likewise, a subset of commonly used KEGG pathways was displayed with a waterfall plot.

### Time series differential expression analysis

Potential contaminants were removed and label free quantification values were log_2_ transformed in Perseus version 1.6.14.0. Data from ESCs and TSCs was processed independently. For both cell type, rows were filtered and any protein which was not expressed in two out of three samples of at least one time point was removed. Missing values were replaced from the normal distribution. Values were extracted and a one-way repeated measures ANOVA with MetaboAnalyst 4.0 was used for time series analysis^67^. Proteins with an adjusted p-value of <0.05 were regarded as differentially expressed.

#### Dynamics of differentially expressed proteins

k-means clustering was used to identify the dynamic behavior of the differentially expressed proteins. The R stats package “stats” (version 4.1.0) was used for k-means clustering.

#### GO term analysis

To identify enriched Biological Processes, Gene Ontology analysis in the clusterProfiler R package (version 4.0.5) was applied on the differentially expressed proteins identified by MetaboAnalyst 4.0. The Benjamini-Hochberg correction was used to correct for multiple comparisons and a pvalueCutoff of 0.05 and qvalueCutoff of 0.1 were used. Enriched biological processes were displayed with dotplot. Selected biological processes were displayed with ggplot2. A full overview of the enriched biological processes is provided in Data S3.

### Metabolomics

ES metabolomics sample prep: 1×108E14 ESCs were collected for metabolomics analysis. Three biological replicates of normal (serum/LIF) and paused (7 days) cells were used.

#### ES metabolomics data acquisition

Metabolite extraction and tandem LC-MS/ MS measurements were done as previously reported by us^68,69^. In brief, methyl-tert-butyl ester (MTBE, Sigma-Aldrich), methanol, ammonium acetate, and water were used for metabolite extraction. Subsequent separation was performed on an LC instrument (1290 series UHPLC; Agilent, Santa Clara, CA), online coupled to a triple quadrupole hybrid ion trap mass spectrometer QTrap 6500 (Sciex, Foster City, CA), as reported previously^70^.

#### ES metabolomics data analysis

The metabolite identification was based on three levels: (i) the correct retention time, (ii) up to three MRM’s (iii) and a matching MRM ion ratio of tuned pure metabolites as a reference. Relative quantification was performed using MultiQuantTM software v. 2.1.1 (Sciex, Foster City, CA), all peaks were reviewed manually. Only the average peak area of the first transition was used for calculations. Normalization was based on cell number of the samples and subsequently by internal standards. The processed output files can be found in Data S4, showing metabolite identification, HMDB and KEGG IDs, log2FC and p-values. Any metabolite with a p-value <0.05 and absolute log2FC >0.75 was regarded as statistically significant. Differentially expressed metabolites were plotted in R with the pheatmap package (version 1.0.12).

#### Embryo metabolomics sample prep

Embryos were cultured in reduced KSOM. The different conditions were identified by fluorescent labeling prior to desiccation. For this, E4.5, D8 mTORi and D15 mTORi + carnitine embryos were stained with MitoSpy Green, MitoSpy Red, and DAPI, respectively. For MitoSpy staining, 800 nM Mito Spy i n r educed KSOM were equilibrated overnight at 5% O2 and 5% CO2 at 37°C. DAPI was dissolved to 300 uM in PBS and equilibrated overnight at 5% O2 and 5% CO2 at 37°C. The embryos were fluorescently labeled by incubating them for 45 minutes in the pre-equilibrated MitoSpy at 5% O2 and 5% CO2 at 37°C or 15 minutes at room temperature in pre-equilibrated DAPI. After staining, the embryos were washed three times with PBS, and then three times with 100 mM ammonium acetate (Merck, cat. # 1.01116.1000). Embryos were deposited in single spots on a microscope slide and desiccated for one hour in a vacuum chamber. Desiccated samples were stored at -80°C until further analysis.

#### Embryo metabolomics data acquisition

Sample slides were removed from -80°C and set in a vacuum chamber for dessication for 1 hour. To avoid condensation, slides were transported directly to the dessicator on dry ice. Matrix deposition was performed using a TM Sprayer (HTX Technologies, Chapel Hill, NC, USA) according to the protocol as follows. Matrix for negative mode was DAN (1,5-Diaminonaphthalene, Sigma) at 7mg/ml concentration sprayed at a constant rate of 0.05ml/min for a total of 7 passes. For positive mode DHB (2,5-Dihydroxybenzoic acid, Sigma) at 15mg/ml concentration sprayed at a constant rate of 0.07ml/min for a total of 8 passes. Mass Spectrometry Imaging was performed with the AP-SMALDI5-Orbitrap MS. The mass range was 100-500 for negative mode 250-1200 for positive mode. After matrix coating, samples were imaged on an AP-SMALDI5 ion source (TransMIT, Giessen, Germany) coupled to a Q Exactive Plus mass spectrometer (Thermo Fisher Scientific, Bremen, Germany). Samples were imaged at 15uM step size with attenuator level 28, to achieve the best spatial resolution in respect to sensitivity and time. Typical acquisition times were around 1-2 hours per sample, sampling around 50 to 100 pixels squared. Thermo RAW data was analyzed by converting to IMZML and IBD format using Thermo imagequest software. Centroiding was performed. IMZML and IBD files were then uploaded to metaspace for result annotation and data interpretation (https://metaspace2020.eu/).

#### Embryo metabolomics data analysis

26 ions with an FDR<10% and embryonic localisation were detected. Metabolite ion intensity was represented as a gray value in grayscale images. To confidently quantify each metabolite across the full embryo respectively, Z projections of all metabolites per spot were created and the embryos masked manually in ImageJ. The resulting masks were then used to quantify the average intensity for each metabolite in its corresponding image. Subsequently, the integrated intensity was calculated by multiplying the average intensity by the number of pixels per embryo. A log_2_FC over E4.5 was computed by calculating the quantified integrated intensity/mean E4.5 integrated intensity per metabolite. The intensity/ average + 0. 01 was log_2_ transformed and mean centered data was visualized with a heatmap. Log_2_FC were plotted using ggplot2 (version 3.3.5).

### Seahorse analysis

#### Seahorse experiment

On the day prior to the assay, sensor cartridges were hydrated with sterile water added to the utility plate of the sensor cartridge and incubated at 37°C in a non-CO2 incubator overnight. 35,000 ES/TSCs were plated onto gelatin coated cell culture miniplates in respective cell culture media and incubated overnight at 37°C and 5% CO2. The next day, the sterile water in the utility plate was replaced with pre-calibrated XF calibrant to calibrate the utility plate of the sensor cartridge for 1 hour at 37°C. The cell culture miniplate having at least 50-60% confluent cells were washed once with 37°C pre-warmed assay medium (XF base medium with 1mM pyruvate, 2mM glutamine and 10mM glucose pH 7.4 and LIF (for ESCs) – mitostress medium or XF base medium with 2mM glutamine pH 7.4 and LIF (for ESCs)

– glycostress medium). After the wash 180ul of new assay medium was added to the cell culture miniplate and the plate was kept at 37°C for 1 hour. After 1 hour of incubation of the sensor cartridge, for mitostress test – components were added in the following order – portA-20ul oligomycin (2uM) ; port B – 22ul FCCP (1uM) ; port C – 25ul Rotenone A/ Antimycin A (0.5uM) and port D – 27ul mitostress assay medium. For glycostress test – components were added in the following order – port A – 20ul glucose (10mM); port B – 22ul oligomycin (2uM) ; port C – 25ul 2-DG (50mM) ; port D – 27ul glycostress assay medium. The sensor cartridge with the utility plate was placed in the Agilent Seahorse XFp Extracellular Flux Analyzer, which was turned on and pre-warmed for 30min for calibration. After the calibration, the utility plate of the sensor cartridge was replaced with a cell culture plate. The Mitostress and Glyostress programs were run using the wave software.

#### Data analysis

Basal respiration and glycolysis rates were extracted. Three replicates were included for each of the two independent biological replicates. The mean and standard error of the mean was calculated based on all six replicates. The basal respiration was plotted against the glycolysis rate with ggplot2 (version 3.3.5).

### Electron microscopy

Diapaused (EDG7.5) or E4.5 embryos were isolated and immediately fixed in 2.5% glutaraldehyde (Grade I, Sigma, Germany) in PBS for 90 minutes at room temperature. The fixed embryos were subsequently embedded in 2% low melting point agarose (Biozym, Hessisch Oldendorf, Germany). Agarose was then carefully cut into cubes with an edge length of roughly 1 mm, containing one embryo inside of each cube. The cubes were collected in 3 ml glass vials and stored overnight at 4°C in PBS. On the following day, samples were post-fixed in 0.5% OsO4 for 2.5 hours at RT on a specimen rotator followed by four washing steps in ddH2O at RT, 5 minutes each. Samples were then incubated for one hour in 0.1% tannic acid (Science Services, Munich, Germany) dissolved in 100mM HEPES on a specimen rotator followed by three washing steps in ddH2O at RT, 10 minutes each. Subsequently, samples were incubated in 2% uranyl acetate (Sigma-Aldrich, Merck, Darmstadt, Germany) for 90 minutes at RT on a specimen rotator. Samples were then washed once in distilled water followed by dehydration through a series of increasing ethanol concentrations (30 min each in 70 %, 90 % and 95 % and finally 3 times 10 min in absolute ethanol, respectively). The dehydrated samples were then incubated in a 1:1 mixture of propylene oxide and absolute ethanol for five minutes at RT followed by a two times 10 minutes incubation in pure propylene oxide (Sigma-Aldrich, Merck, Darmstadt, Germany), and a 30 minutes incubation in a 1:1 mixture of propylene oxide and Spurr resin (Low Viscosity Spurr Kit, Ted Pella, Plano Gmb H, Wetzlar, Germany) at RT. Specimens were finally transferred to pure Spurr resin mixture and infiltrated overnight at 4°C. Spurr mixture was renewed for two times 2 hours on the following day. Afterwards resin infiltrated embryos in agarose cubes were polymerized at 60°C for forty-eight hours. Ultrathin sections (70nm) were cut from polymerized samples using a Leica UC7 ultramicrotome equipped with a 3 mm diamond knife (Diatome, Biel, Switzerland) and placed on 3.05 mm Formvar Carbon coated TEM copper slot grids (Plano GmbH, Wetzlar, Germany). Sections were post contrasted with UranyLess EM stain and lead citrate (both from Science Services, Munich, Germany). In order to visualize ultrastructural details, sections were imaged fully automatically at 4400x nominal magnification using Leginon (Suloway et al., 2005) on a Tecnai Spirit transmission electron microscope (FEI) operated at 120 kV which was equipped with a 4kx4k F416 CMOS camera (TVIPS). Acquired images were then stitched to a single montage using the TrakEM2 plugin^71^ implemented in Fiji^72^.

#### Analysis

Analysis of the electron microscopy images was carried out semi-automated using the ZEN 3.4 Blue software (Zeiss). The blastocyst structures together with the analyzed features (lipid droplets, mitochondria and nuclei) were manually masked as primary objects and added to the analysis and measurements (area, feret ratio, roundness, etc.) were selected, respectively. The ZEN 3.4 Blue inbuilt function Zone of Influence (ZOI) analysis was carried out on the masked images to determine the number of mitochondria surrounding each lipid droplet (radius was set to 80 pixel). In the cases where lipid droplets occurred as clusters, binary dilation with a count of 15 was used to fuse the lipid droplets in a cluster and these clusters were analyzed as one droplet to prevent oversampling.

### Embryo stainings

#### Experimental setup

Embryos were cultured in reduced KSOM, paused via mTOR inhibition with 200 nM Rapalink and additionally supplemented with 1 mM L-carnitine. Embryos were collected at E4.5, D3 mTORi +/- L-carnitine, D8 mTORi +/- L-carnitine and D15 mTORi + carnitine. Additionally, D8 mTORi +/- L-carnitine and D15 mTORi + carnitine were allowed to reactivate for 12 or 24 hours prior to staining.

#### Embryo immunofluorescence stainings

Embryos were collected and fixed for 10 minutes in 4% PFA, then permeabilized for 15 minutes in 0.2% Triton X-100 (Sigma, cat. # T8787) in PBS. Permeabilized embryos were incubated in blocking buffer (0.2% Triton X-100 in PBS + 2% BSA (BSA Fraction V 7.5%, Gibco, cat.# 15260-037) + 5% goat serum (Jackson Immunoresrearch/ Dianova, cat. # 017-000-121) for 1h. Then embryos were incubated overnight at 4°C with the primary antibodies. Primary antibodies were diluted in blocking buffer, 1:50 for the Oct3/4 mouse anti-mouse (Santa Cruz, cat.# sc-5279), 1:100 for the H4K16ac rabbit anti-mouse antibody (Millipore, cat.# 7329), 1:100 for the c-Caspase3 rabbit anti-mouse antibody (Cell Signaling Technology, cat.# 9661S), 1:50 for the FoxO1 rabbit anti-mouse antibody (Cell Signaling Technology, cat.# 2880T), 1:100 for the Anti-Lamin B1 antibody - Nuclear Envelope Marker (Abcam, cat#. 16048), and 1:100 for the Anti-Histone H3 (di methyl K9) antibody (Abcam, cat.# 1220). Prior to incubation with the secondary antibodies, embryos were washed three times in washing buffer (0.2% Triton X-100 in PBS + 2% BSA).

Then, the embryos were incubated with the secondary antibodies for 1h at room temperature. The donkey anti-rabbit Alexa Fluor Plus 647 (Themo, cat.# A32795) and donkey anti-mouse Alexa Fluor 488 (Thermo, cat.# A21202) antibodies were diluted 1:200 in blocking buffer. After incubation with the secondary antibody, embryos were washed three times in washing buffer. Then, embryos were mounted on a microscope slide with a Secure-Seal™ Spacer (8 wells, 9 mm diameter, 0.12 mm deep, ThemoFisher, cat.# S24737), covered with a cover glass and sealed with nail polish.

#### Bodipy stainings

Embryos were stained for 30 minutes at 5% O2 and 5% CO2 at 37°C with 11.4 µM Bodipy, 5 µM CellMask Deep Red (ThermoFisher, cat.# A57245), and 16.2 µM Hoechst 33342 (ThermoFisher, cat.# H3570) in PBS. Stained embryos were washed with PBS and directly imaged in PBS on an Ibidi µ-Slide 8 Well (Ibidi, cat.# 80821).

#### Imaging

Embryos were imaged with the Zeiss Plan-Apochromat 20x/0.8 objective on the Zeiss LSM880 Airy microscope using Airy scan. Airyprocessing (2D, strength of 1) was done with the Zen Black software. Images were further processed using Fiji ImageJ2 (version 2.3.0). Image quantifications were done with CellProfiler (version 4.2.1). For the H4K16ac and OCT4 qualifications, nuclei were identified as primary objects using the DAPI stain. The integrated intensities of OCT4 and H4K16ac were normalized to the DAPI intensity. For the FOXO1 stainings, first nuclei were identified as primary objects, then cellular boundary was set by expanding the primary object until touching the next cell. The integrated intensity of FOXO1 was normalized to the cellular area. For the Bodipy stainings, the number of lipid droplets per embryo were quantified by identifying the embryo as primary object and the LDs as objects. Only LDs in the embryo were counted. For the per cell quantification, nuclei were identified as primary objects and enlarged until touching to identify cells. The cytoplasm was derived by subtracting the nuclear area from the cells. Additionally, LDs were identified as primary objects as only those identified in the cytoplasm were used for further analysis.

---

**Figure S1.**
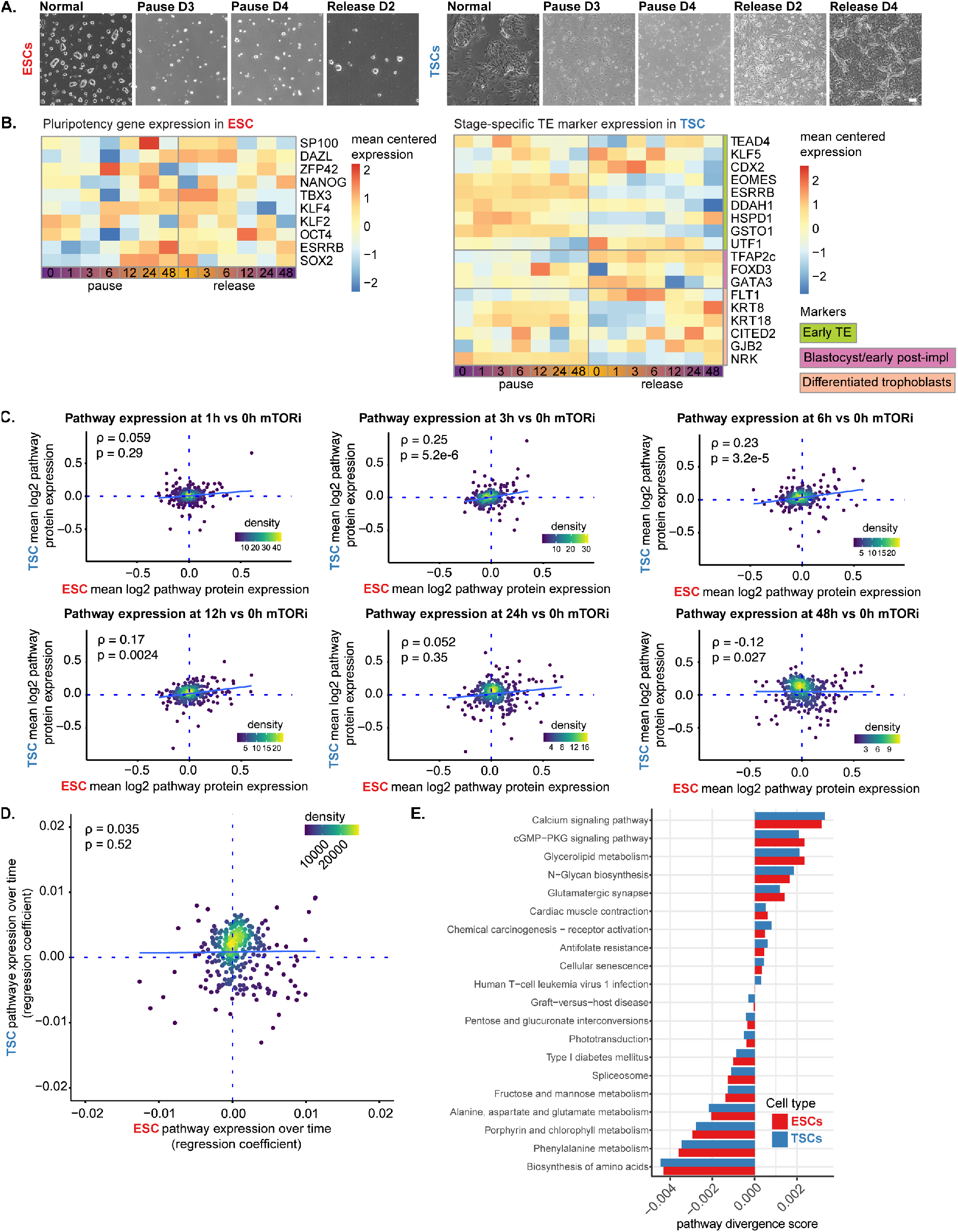
Pathway expression over time and commonly used pathways. **(A)** Bright field images of ESCs and TSCs during normal proliferation, at D3 and D4 of pausing and upon release from mTORi. Scale bar: 100 µm. **(B)** Mean protein expression (pause and release vs control t=0) of pluripotency genes in ESCs and stage-specific TE marker expression in TSCs (previously curated gene list^73^. **(C)** Log2FC of mean protein expression (mTORi/control t=0) of each KEGG pathway at each time point. **(D)** Change in pathway expression over time, computed by a linear regression of the mean KEGG pathway protein expression. **(E)** Commonly used KEGG pathways in ESCs and TSCs in response to mTORi.

**Figure S2.**
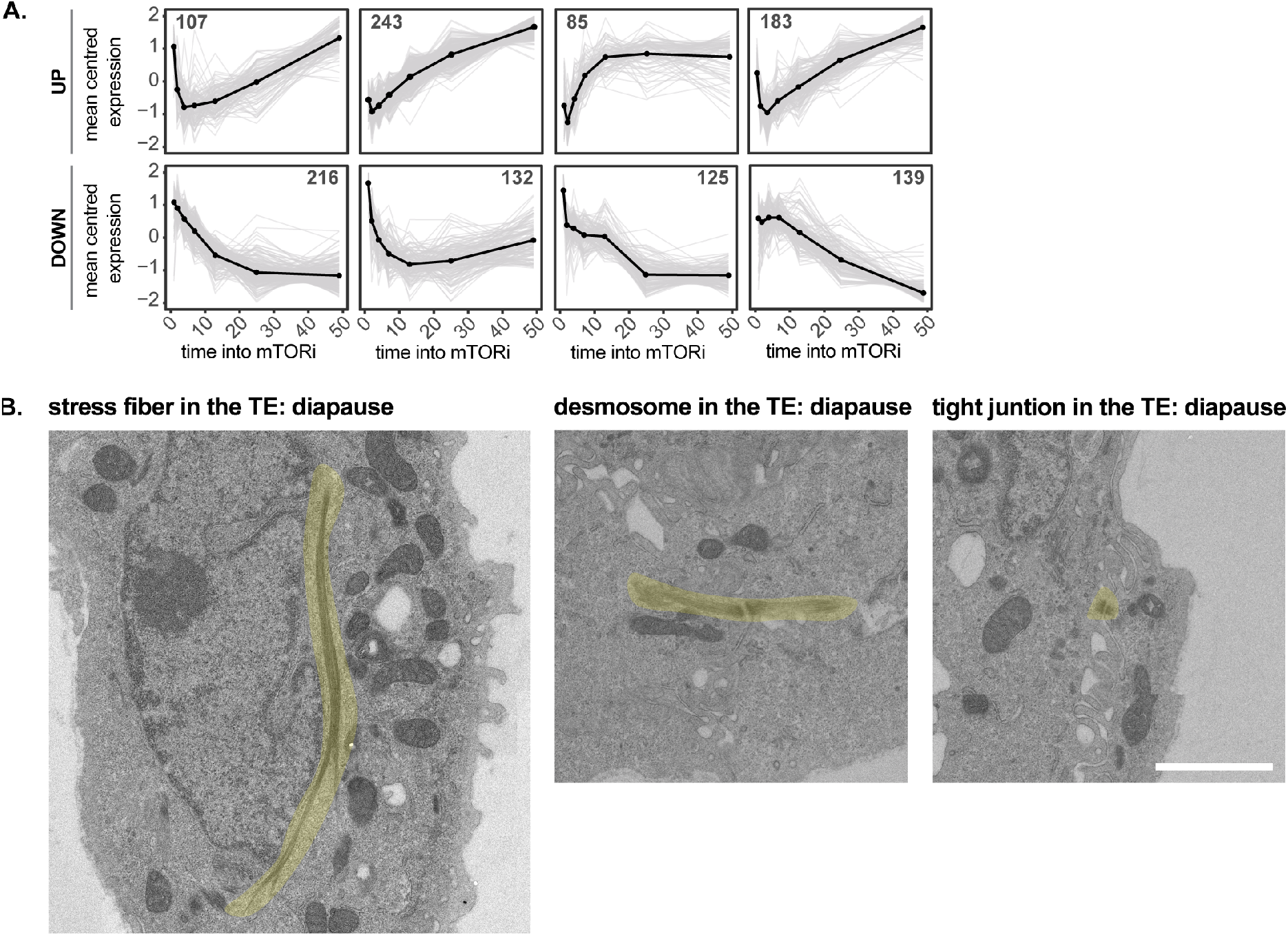
Proteome rewiring during entry into pausing in TSCs. **(A)** K-means clustering of differentially expressed proteins identified with MetaboAnalyst 4.0. **(B)** Transmission electron microscopy images of cell-cell adhesion complexes and stress fibers. Structures are highlighted in yellow. Scale bar: 2 µm.

**Figure S3.**
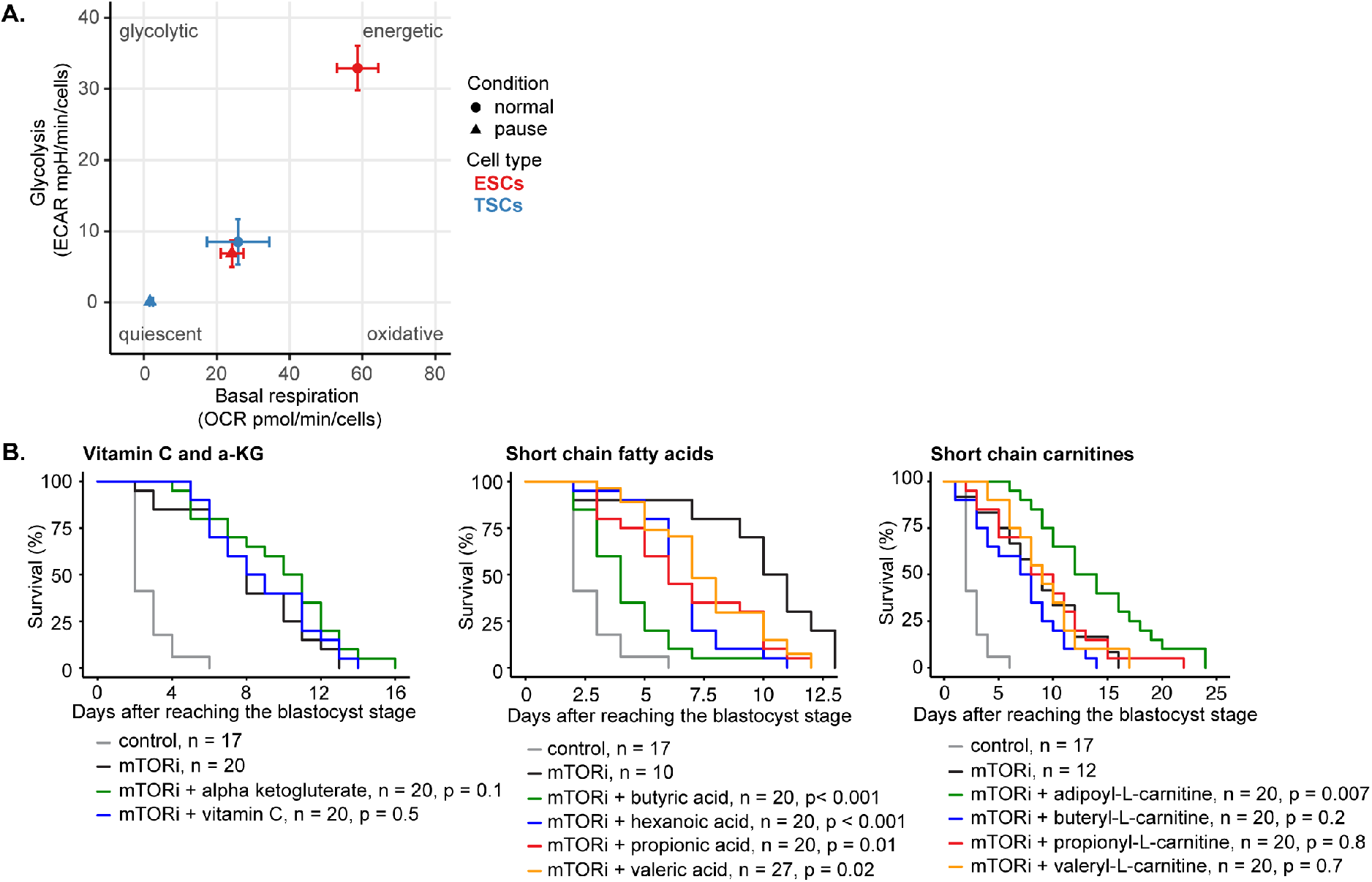
Energetics in ESCs and TSCs, and embryo supplement survival curves. **(A)** Energetics of normal and paused ESCs and TSCs. The mean +/- SEM of three replicates of two independent biological replicates are shown. **(B)** Survival curves or paused embryos supplemented with vitamin C, alpha-ketoglutarate, short chain fatty acids, and short chain carnitines. n = number of embryos. Statistical test is the G-rho family test of Harrington and Fleming.

**Figure S4.**
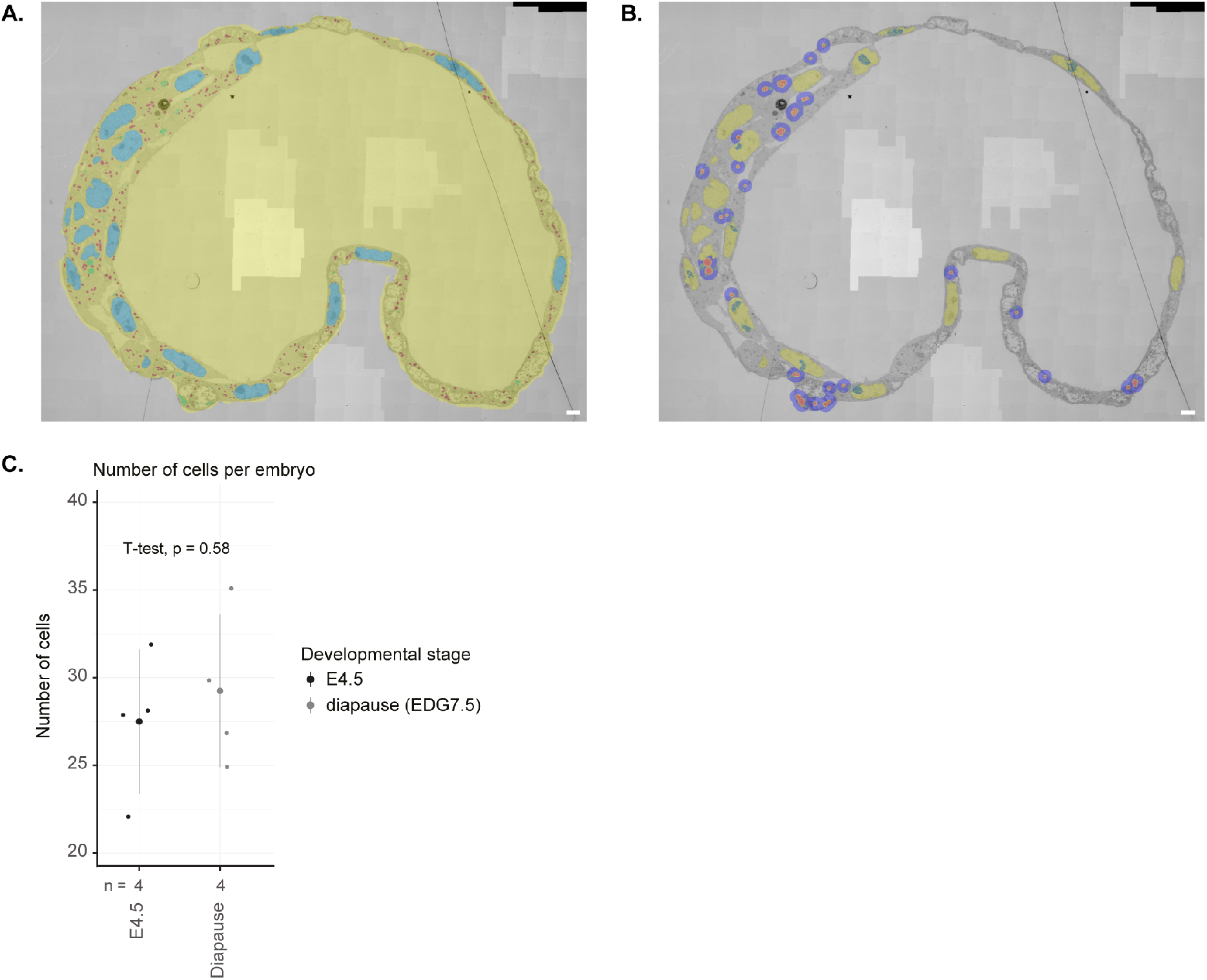
LD and mitochondria quantifications of EM images. **(A)** Example masking of the embryo (yellow), nucleus (light blue), LDs (green), and mitochondria (red). Scale bar: 2 µm. **(B)** Masking of the zone of influence to identify the number of mitochondria in close proximity to each LD or a cluster of LDs. Masked are the nucleus (yellow), the LDs (red), a watershed around the lipid droplets (orange), the zone of interest of 80 pixels around the watershed object containing the lipid droplets (purple), and the mitochondria in the zone of interest (green). Scale bar: 2 µm. **(C)** Number of analyzed cells per embryo, as quantified using the EM images. n = number of embryos per condition.

**Figure S5.**
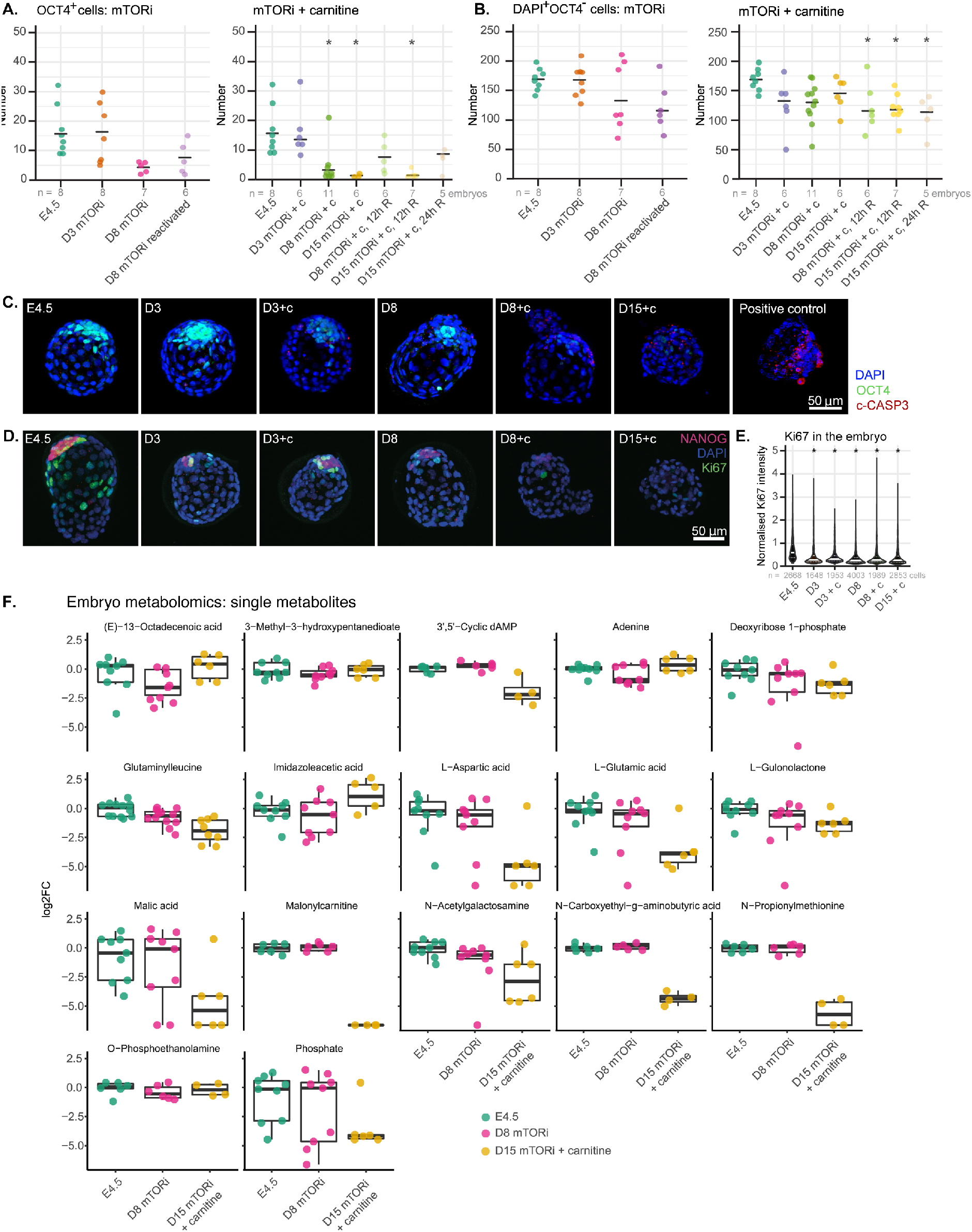
Cell number dynamics, apoptosis, and single embryo metabolomics data. **(A)** Number of OCT4-positive cells per embryo in each condition. n = number of embryos, *: p <0.05, one way ANOVA, Dunnett’s post hoc test. **(B)** Number of TE cells per embryo in each condition. TE cells are defined as DAPI-positive and OCT4-negative. n = number of embryos, *: p <0.05, one way ANOVA, Dunnett’s post hoc test. **(C)** Representative IF images of embryos stained against cleaved-CASPASE3 and OCT4 in each condition (n = 5 embryos/condition). As positive control, E4.5 embryos were exposed to UV with a UV crosslinker. energy 4000 uJ/cm2 for 8 seconds and subsequently cultured for another 6 hours. Across conditions, some TE cells display cleaved-caspase3 staining. Scale bar = 50 µm. **(D)** Representative IF images of embryos stained against Ki67 and NANOG in each condition (n = 8 embryos/ condition). Scale bar = 50 µm.

**Figure S6.**
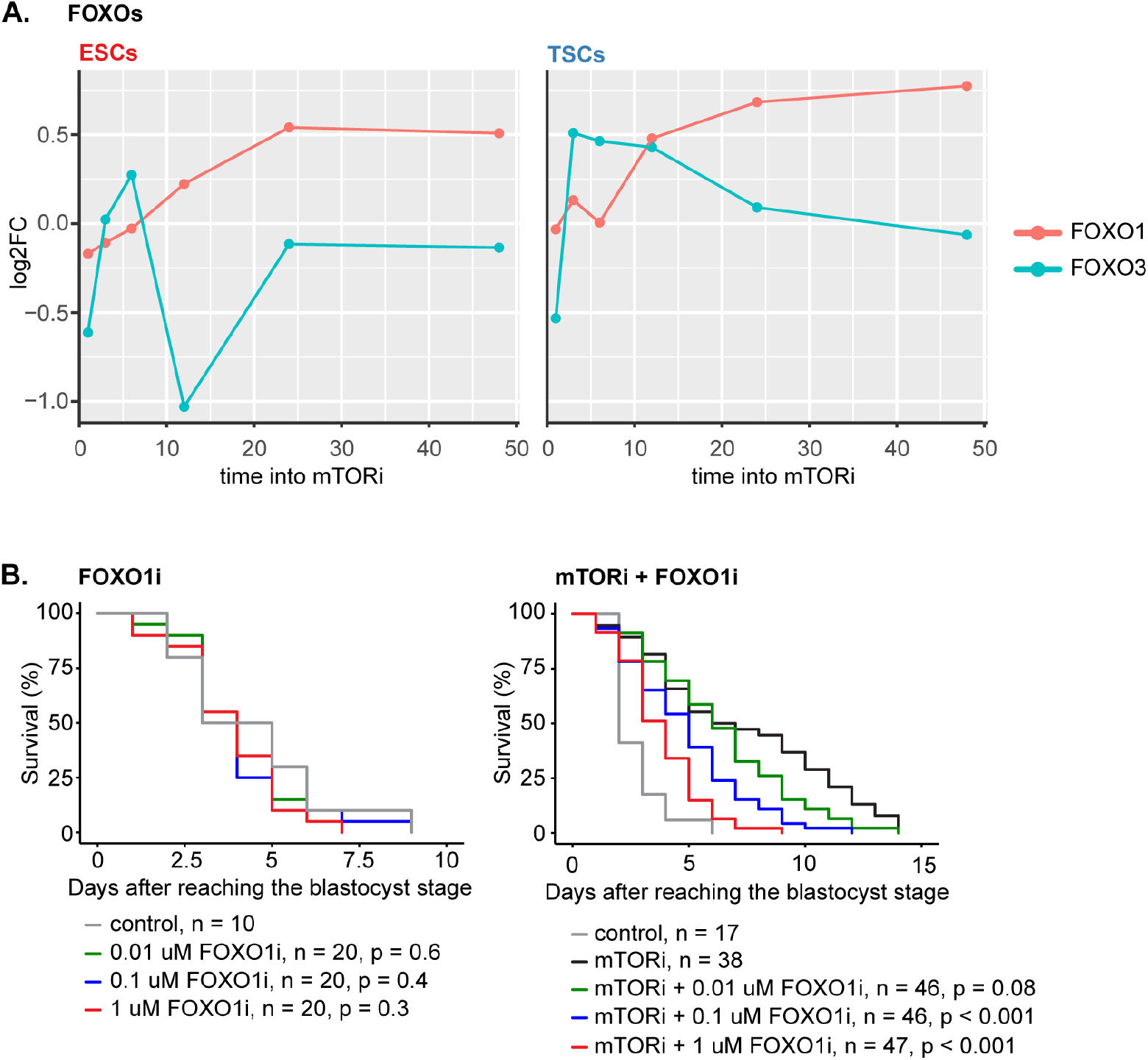
FOXO1 and FOXO3 expression data and FOXO1i survival curves. **(A)** FOXO1 and FOXO3 protein expression into mTORi in ESCs and TSCs. **(B)** Embryo survival curves in the shown conditions.

